# What mycologists should talk about when they are talking about the International Code of Nomenclature for algae, fungi, and plants

**DOI:** 10.1101/2023.02.25.529721

**Authors:** R. Henrik Nilsson, Martin Ryberg, Christian Wurzbacher, Leho Tedersoo, Sten Anslan, Sergei Põlme, Viacheslav Spirin, Vladimir Mikryukov, Sten Svantesson, Martin Hartmann, Charlotte Lennartsdotter, Pauline Belford, Maryia Khomich, Alice Retter, Natàlia Corcoll, Daniela Gómez Martinez, Tobias Jansson, Masoomeh Ghobad-Nejhad, Duong Vu, Marisol Sanchez-Garcia, Erik Kristiansson, Kessy Abarenkov

## Abstract

Fungal metabarcoding of substrates such as soil, wood, and water are uncovering an unprecedented number of fungal species that do not seem to produce tangible morphological structures and that defy our best attempts at cultivation, thus falling outside of the ambit of the International Code of Nomenclature for algae, fungi, and plants. The present study uses the new, ninth release of the species hypotheses of the UNITE database to show that species discovery through environmental sequencing vastly outpaces traditional, Sanger sequencing-based efforts in a strongly increasing trend over the last five years. Our findings challenge the present stance of the mycological community – that “the code” works fine and that these complications will somehow sort themselves out given enough time and a following wind – and suggest that we should be discussing not whether to allow DNA-based descriptions (typifications) of species and by extension higher ranks of fungi, but what the precise requirements for such DNA-based typifications should be. We submit a tentative list of such criteria for further discussion. However, the present authors fear that no waves of change will be lapping the shores of mycology for the foreseeable future, leaving the overwhelming majority of extant fungi without formal names and thus scientific and environmental agency. It is not clear to us who benefits from that, but neither fungi nor mycology are likely to be on the winning side.

## Introduction

Dark matter is an astronomical concept that denotes mass of a hitherto unknown nature. That mass is detectable indirectly through the gravity it exerts – such as the bending of passing light – but its exact nature has so far defied scientific explanation. Mycology offers an analogy in the form of dark taxa, which are taxa that do not seem to produce tangible morphological structures and that we cannot seem to cultivate in the lab. As with dark matter, dark taxa are chiefly detected by other means than direct observation, notably through DNA sequencing (Grossart et al. 2016; Lücking et al. 2021). The field of mycology has become intimately entwined with the concept of dark taxa in the wake of environmental metabarcoding, where seemingly dark taxa often make up more than half of the taxa recovered (e.g., Retter et al. 2019). Dark taxa seem to permeate the fungal tree of life and are known from all major fungal lineages. Indeed, a non-trivial number of large fungal lineages are constituted solely by dark taxa (Tedersoo et al. 2017, 2020). Studying the fungal kingdom sans its dark components is to study a paraphyletic group, something that contemporary phylogenetic thinking advises strongly against.

Most of the present authors have spent considerable time in the company of dark fungal taxa (DFT) as recovered through environmental metabarcoding and as manifested in the UNITE database for molecular identification of fungi (Nilsson et al. 2019). The sheer magnitude of extant sequence data from DFT signals a need to take these taxa seriously. Yet it seems to the present authors that contemporary mycology treats DFT as if they had a lesser in fact, no – biological validity. The International Code of Nomenclature for algae, fungi, and plants (ICN; Turland et al. 2018) does not permit species descriptions typified from DNA sequences alone, and a recent effort to bring about change in this regard was overthrown with overwhelming majority (May et al. 2018). Similarly, DFT are routinely ignored in the context of, e.g., phylogenetic inference, ecology, and nature conservation (Ryberg and Nilsson, 2018). Indeed, it is as if the DFT have no agency at all, scientific or otherwise. This goes very much against the experience of the present authors, who have used DFT to tease out branching orders, dominant but entirely overlooked taxa, and major ecological patterns that otherwise would have been lost on science (Khan et al. 2020; Nilsson et al. 2011, 2016; Tedersoo et al. 2022). Similarly, in an attempt to accord some taxonomic standing to the DFT, UNITE has assigned DOI-based digital identifiers to all DFT known from nuclear ribosomal internal transcribed spacer region (ITS) data to facilitate and promote unambiguous scientific communication across datasets and studies (Kõljalg et al. 2013). These efforts have largely fallen short of sparking the debate they were hoping to.

In the present forum paper, we wish to visualize the relative contribution of DFT to molecular mycological species discovery over time. We do this through two molecular datasets, both of which reflect current knowledge but also biases in various ways. These datasets are: 1) all full-length fungal ITS sequences in the international sequence database collaboration (INSDC; Arita et al. 2021) as of October 11, 2022 and 2) the five large metabarcoding datasets – chiefly of soil fungi (e.g., Tedersoo et al. 2022) – so far incorporated into the UNITE database. We find that the DFT overwhelmingly dominate the species discovery process, and it seems patently clear that extant fungal diversity presents us with patterns that cannot be accurately represented only by species defined by morphology or cultivation alone. It strikes us as unfortunate that what seems to be the absolute majority of fungi fall outside the ambit of the ICN, and we hope that the present results will instigate a much needed – and much overdue – debate on how and when we should allow formal species descriptions based on DNA sequence data alone.

## Materials and Methods

The full flow of operation behind the UNITE database is described elsewhere (Kõljalg et al. 2013, 2020; Nilsson et al. 2019). In brief, UNITE clusters the fungal ITS sequences of INSDC jointly with the UNITE-contributed DFT ITS sequences into species hypotheses (SHs) at distance thresholds 0.5% through to 3.0% in steps of 0.5%. These operational taxonomic units can be thought of as entities roughly at the species level. The sequences and the SHs are available for web-based interaction as well as for download in various formats (https://unite.ut.ee/repository.php).

We downloaded all sequences included in the October 2022 version 9 release of the UNITE species hypothesis system. To allow us to contrast the species discovery from taxonomic and metabarcoding studies, we made the admittedly coarse assumption that all SHs that contained at least one sequence from the INSDC could be considered as taxonomy-derived SHs, that is, SHs with some sort of footing in traditional taxonomy. In analogy, all SHs comprised solely of metabarcoding sequences were considered as DFT. Based on the date of initial submission of each sequence (submission to INSDC and to UNITE, respectively, for INSDC and DFT sequences), we examined the accumulation of SHs over time. We plotted the accumulation of taxonomy-derived and DFT-only SHs against date of initial discovery in R v. 4.2.2 (R core team 2020).

While there is little hope of piecing together the ecological context of these sequences in an automated way, at least there is an opportunity to visualize the country of collection for many of the sequences in INSDC and UNITE. We thus sought to illustrate the geographical component of the SH accumulation curves by summarizing the country of collection of the taxonomy-derived and DFT sequences. In total, 63% of the taxonomy-derived, and 99.9% of the DFT, sequences were tagged with an explicit country of origin. The 20 most common countries of origin in each dataset were compiled using R.

## Results

We retrieved a total of 1.26 M taxonomy-derived sequences from INSDC and 7.1 M DFT sequences from UNITE. The taxonomy-derived sequences were found to stem from a total of 88,665 distinct published and unpublished studies as defined by the combination of the INSDC fields AUTHORS, TITLE, and JOURNAL. The DFT sequences were found to stem from 5 studies. The SH accumulation curves at the dynamic 1.5% similarity threshold level are shown in Figure 1. Table 1 shows the top 20 countries of origin for the taxonomy-derived and DFT sequences for which this data was available. Figure 2 shows the collection localities for all Sanger and metabarcoding sequences with geo-coordinates.

**Table 1.** The 20 most common countries of collection for the Sanger and the DFT sequences. The DFT dataset is dominated by sequences from Estonia, from which the five metabarcoding studies were run. Estonia is not known as any particularly rich hotspot of biodiversity, perhaps suggesting that additional worldwide sampling would have produced even more dramatic increases in the number of DFT SHs.

| 1 | INSDC country | INSDC seq. | DFT country | DFT seq. |
| --- | --- | --- | --- | --- |
| 2 | Unknown | 463524 | Estonia | 1788894 |
| 3 | United States | 133496 | United States | 350869 |
| 4 | China | 117292 | Italy | 287842 |
| 5 | India | 31788 | Brazil | 285473 |
| 6 | Japan | 29754 | Czechia | 260611 |
| 7 | Brazil | 27765 | Russian Federation | 228979 |
| 8 | Canada | 26038 | Mexico | 210643 |
| 9 | Spain | 22362 | Norway | 208422 |
| 10 | Australia | 22205 | Colombia | 204172 |
| 11 | Germany | 19971 | Australia | 177777 |
| 12 | Italy | 18078 | Sweden | 177318 |
| 13 | Mexico | 16326 | Latvia | 169168 |
| 14 | France | 14896 | Lithuania | 166553 |
| 15 | Korea, Republic of | 12434 | Georgia | 146440 |
| 16 | Russian Federation | 11668 | Finland | 127258 |
| 17 | Iran, Islamic Republic of | 11285 | India | 123706 |
| 18 | Poland | 10969 | Argentina | 116852 |
| 19 | New Zealand | 10956 | China | 100143 |
| 20 | Thailand | 10708 | Papua New Guinea | 96253 |
| 21 | South Africa | 10642 | Tanzania, United | 95203 |

**Figure 1.**
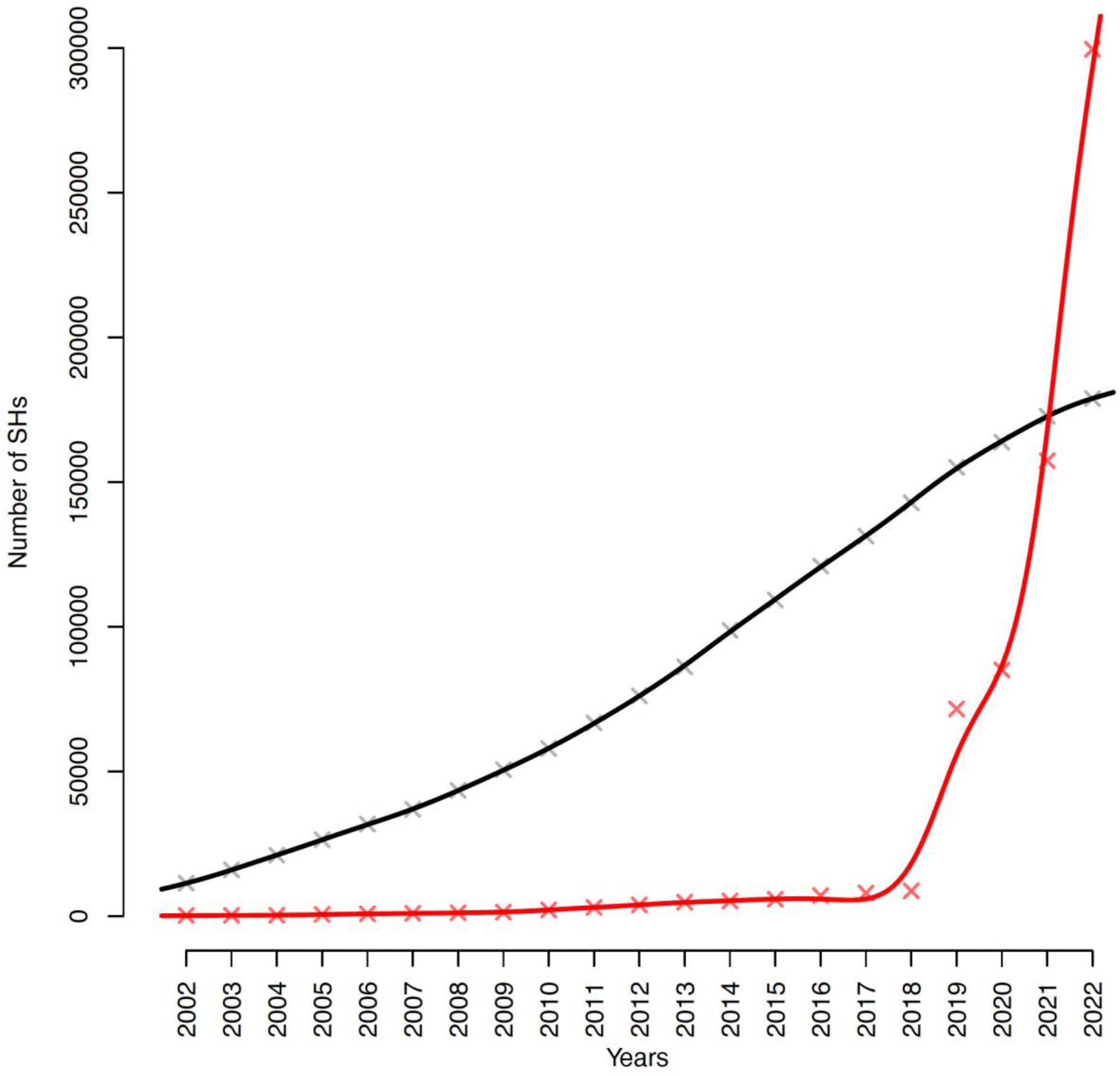
The accumulation of SHs at the 1.5% distance threshold over time in the Sanger (black; 88,665 studies of various sizes) and the DFT (red; 5 large studies) datasets. The Y axis depicts the number of SHs and the X axis depicts year of sequence deposition. Solid trend lines were calculated using cubic smoothing splines.

**Figure 2.**
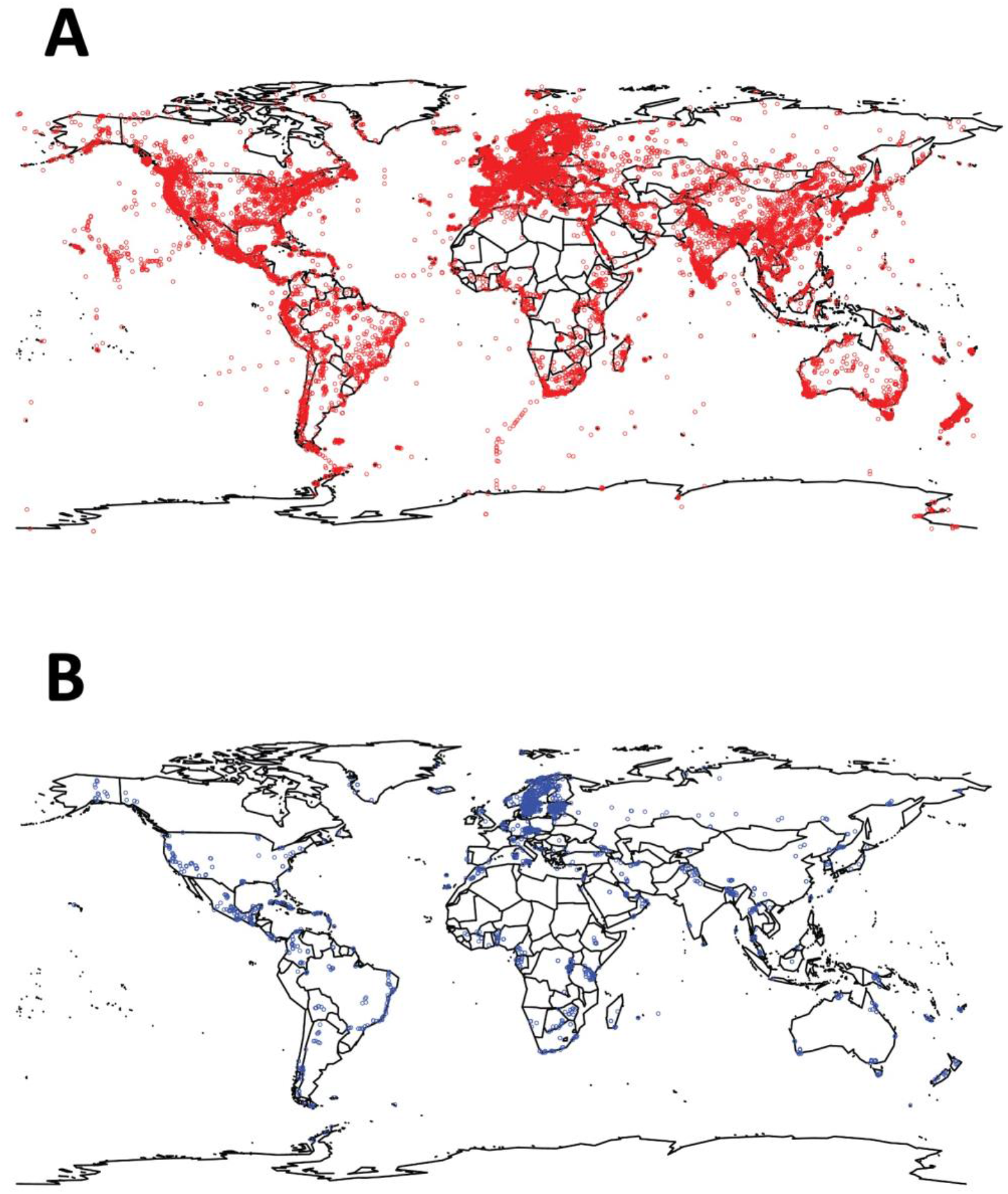
Maps showing the collection localities for the (A) Sanger sequences and (B) metabarcoding sequences that came with geo-coordinates (36,559 Sanger collection localities and 3,688 metabarcoding collection localities).

## Discussion

The present study approximated fungal species accumulation over time as deduced from taxonomic and metabarcoding efforts. We found that the DFT account for the lion’s share of the new species discovered in the last five years (although some limited proportion of both the Sanger-derived and the DFT sequences may possibly correspond to described, but so far unsequenced, species). We reached this conclusion based on a very limited number of studies in fact, just five – on soil fungal communities and in almost complete absence of metabarcoding data from, e.g., water, air, wood, and plant material. One can only imagine that Figure 1 would have shown an even more dramatic trend had a wider selection of metabarcoding datasets been available in UNITE. Figure 2 paints a similar picture with respect to the geographical coverage. It shows that whereas the sampling effort of the five metabarcoding studies was wide, it pales in comparison to that of the combined Sanger-derived studies. It is reasonable to think that at least some of the unsampled geographical regions are rich in DFT and would have contributed to an even steeper trend in Figure 1, had they been sampled.

It is often said that when data are sparse, opinions may be maintained and cherished for longer than necessary. Our results show that data are no longer sparse; DFT, in view of their diversity and abundance, form a major, inextricable component of the fungal kingdom. They simply cannot be swept under the carpet. It is not scientifically defensible to exclude them from mycological efforts in phylogeny, ecology, or biogeography; they simply cannot be swept under the carpet. We therefore argue that it does not make sense to deny them a formal standing under the ICN. It is time – in fact, long overdue – to start discussing what the requirements should be for DFT to be formally considered under the ICN. Clearly, morphological structures or cultivability cannot be part of those requirements. We feel it is time for serious discussion on this topic, and we would like to reiterate the observations of Lücking et al. (2021) that a limited number of thought-through requirements would probably suffice. These should reflect the need for scientific reproducibility and should be stringent enough that only particularly well-vetted and documented DFT can be considered for DNA-based typification and formal description. At the same time, they should be realistic and reasonable enough that formal taxonomic description does become possible for such particularly well-vetted and documented DFT. We submit the following as tentative criteria:

- A minimum length/coverage for the underlying sequence data (such as all of SSU, ITS, and LSU in a contiguous stretch).
- Sufficiently high read quality.
- At least two independent recoveries of the taxon across separate datasets, perhaps from separate research teams.
- A thorough analysis of the public sequence databases for relevant additional sequences to maximise the penetration of available data and to minimize redundant descriptions.
- An underlying phylogenetic analysis based on a multiple sequence (perhaps SSU plus LSU, or at least LSU) alignment.
- Bundling of open, richly annotated raw sequence data/FASTQ/chromatograms and metadata on, e.g., the ecological and geographical specifics of the sampling sites.
- Publication in a scientific journal with a formal impact factor.
- It furthermore seems reasonable to us to allow DNA sequences as types only in fungal groups that are predominantly or exclusively dark at, say, the order or supra-order level. We would be against DNA-based typifications in groups where morphological structures and/or cultivation may be within reach (e.g., *Cortinarius* and *Fusarium*).
- At least one mycological taxonomist should be involved in the description of DFT (indeed, all fungi). There is no shortage of potential complications that, when overlooked, could lead to needless and haphazard introduction of new species and genera in DFT and beyond. For instance, it is well known that some extant genera offer examples of very divergent ITS (or other ribosomal) regions (e.g., *Basidiodendron, Oliveonia*, and *Cantharellus*; Feibelman et al. 1994; Alm Rosenblad et al. 2022). When considered in isolation and out of context – in, say, a molecular ecology dataset – such sequences could be interpreted to warrant new species and genus descriptions. Needless to say, the present authors are against premature description of species and other taxonomic groups.

There is clearly room for refinement of the requirements mentioned here, and we are furthermore certain that the mycological community can come up with additional prerequisites to further increase stringency and reduce the risk for haphazard, more or less irreproducible or irresponsible use of DNA sequences as types (cf. Hibbett et al. 2016; Zamora et al. 2018; Lücking et al. 2018). The present authors warmly welcome – indeed, invite – such a discussion.

It could be argued that a separate nomenclature code should be erected for the DFT, akin perhaps to the *Candidatus* concept in bacteria (Murray and Stackebrandt 1995) or to the extant DOI-based species identifiers of UNITE. We remain sceptical, however, and we argue for full-fledged integration of the DFT into the ICN. After all, the *Candidatus* concept never really took off, and the UNITE DOIs for DFT remain under-used. It seems likely to us that DFT as governed by a separate and more or less unofficial code would simply remain relegated to some state of secondary – in practise, no – importance. That is not the message delivered by Figure 1, however, and not a state fit to reflect the crucial roles fungi are increasingly understood to play in the ecosystems of the world – by the scientific community and the general public alike. On the contrary, DFT seem to dominate the fungal kingdom. This puts the ICN in a position where it governs an ever-dwindling proportion of the extant fungi – maybe just some few percent. Such a position would seem untenable and, ultimately, vulnerable to usurpation. After all, the new and rebellious prokaryotic SeqCode (Hedlund et al. 2022) grew out of frustration at the inability of the International Code of Nomenclature of Prokaryotes (ICNP) to adapt enough to be able to reflect extant prokaryotic diversity properly. While the ultimate fate of SeqCode remains to be seen (Marinov et al. 2022), it does set an eerie example of what the future may hold in store for ICN should DFT continue to be dismissed as irrelevant. We argue that formal scientific names for DFT are necessary for their scientific agency. Similarly, formal names will in practice be needed in biological conservation and in efforts exploring DFT for, e.g., medical and industrial use. These fungi deserve and need formal names, and it is our firm belief and opinion that this is achievable.

## Conclusion

The concept of dark taxa draws from the astronomical concept of dark matter. In the context of the latest release of the UNITE SH system, astronomy offers one further analogy: that of the night sky. Much as stars form tiny specks of light against the massively dark expanse of space, taxonomy-derived SHs in the ninth UNITE SH release give a diminutive impression against the massive backdrop of DFT SHs. It was roughly 100 years ago that astronomy came to terms with the fact that space stretched far beyond our own galaxy, and it is only fit that mycology finally reaches a comparable conclusion with respect to extant fungal diversity.

## Acknowledgements

The GenBank staff is gratefully acknowledged for assistance with establishing the date of submission for the INSDC entries.

## References

Alm Rosenblad M, Larsson E, Walker A, Thongklang N, Wurzbacher C, Nilsson RH (2022) Evidence for further non-coding RNA genes in the fungal rDNA region. MycoKeys 90: 203–213. doi: 10.3897/mycokeys.90.84866

Arita M, Karsch-Mizrachi I, Cochrane G (2021) The international nucleotide sequence database collaboration. Nucleic Acids Research 49: D121–D124. https://doi.org/10.1093/nar/gkaa967

Feibelman T, Bayman P, Cibula WG (1994) Length variation in the internal transcribed spacer of ribosomal DNA in chanterelles. Mycological Research 98(6): 614–618. https://doi.org/10.1016/S0953-7562(09)80407-3

Grossart H-P, Wurzbacher C, James T, Kagami Maiko (2016) Discovery of dark matter fungi in aquatic ecosystems demands a reappraisal of the phylogeny and ecology of zoosporic fungi. Fungal Ecolology 19: 28–38. https://doi.org/10.1016/j.funeco.2015.06.004

Hedlund BP, Chuvochina M, Hugenholtz P, Konstantinidis KT, Murray AE, Palmer M, Parks DH, Probst AJ, Reysenbach A-L, Rodriguez LM, Rossello-Mora R, Sutcliffe IC, Venter SN, Whitman WB (2022) SeqCode: a nomenclatural code for prokaryotes described from sequence data. Nature Microbiology 7(10): 1702–1708. https://doi.org/10.1038/s41564-022-01214-9

Hibbett D, Abarenkov K, Kõljalg U, Öpik M, Chai B, Cole J, Wang Q, Crous P, Robert V, Helgason T, Herr JR, Kirk P, Lueschow S, O’Donnell K, Nilsson RH, Oono R, Schoch C, Smyth C, Walker DM, Porras-Alfaro A, Taylor JW, Geiser GM (2016) Sequence-based classification and identification of Fungi. Mycologia 108(6): 1049–1068. https://doi.org/10.3852/16-130

Khan F, Kluting K, Tångrot J, Urbina H, Ammunet T, Sahraei SE, Rydén M, Ryberg M, Rosling A (2020) Naming the untouchable–environmental sequences and niche partitioning as taxonomical evidence in fungi. IMA Fungus 11: 1–12. https://doi.org/10.1186/s43008-020-00045-9

Kõljalg U, Nilsson RH, Abarenkov K, Tedersoo L, Taylor AFS, Bahram M, Bates ST, Bruns TD, Bengtsson-Palme J, Callaghan TM, Douglas B, Drenkhan T, Eberhardt U, Dueñas M, Grebenc T, Griffith GW, Hartmann M, Kirk PM, Kohout P, Larsson E, Lindahl BD, Lücking R, Martín MP, Matheny PB, Nguyen NH, Niskanen T, Oja J, Peay KG, Peintner U, Peterson M, Põldmaa K, Saag L, Saar I, Schüßler A, Scott JA, Senés C\, Smith ME, Suija A, Taylor DL, Telleria MT, Weiss M, Larsson K-H (2013) Towards a unified paradigm for sequence-based identification of fungi. Molecular Ecology 22(21): 5271–5277. https://doi.org/10.1111/mec.12481

Kõljalg U, Nilsson HR, Schigel D, Tedersoo L, Larsson K-H, May TW, Taylor AFS, Stjernegaard Jeppesen T, Guldberg Frøslev T, Lindahl BD, Põldmaa K, Saar I, Suija A, Savchenko A, Yatsiuk I, Adojaan K, Ivanov F, Piirmann T, Pöhönen R, Zirk A, Abarenkov K (2020) The taxon hypothesis paradigm - On the unambiguous detection and communication of taxa. Microorganisms 8(12): 1910. https://doi.org/10.3390%2Fmicroorganisms8121910

Lücking R, Kirk PM, Hawksworth DL (2018) Sequence-based nomenclature: a reply to Thines et al. and Zamora et al. and provisions for an amended proposal “from the floor” to allow DNA sequences as types of names. IMA Fungus 9: 185–198. https://doi.org/10.5598/imafungus.2018.09.01.12

Lücking R, Aime MC, Robbertse B, Miller AN, Aoki T, Ariyawansa HA, Cardinali G, Crous PW, Druzhinina IS, Geiser DM, Hawksworth DL, Hyde KD, Irinyi L, Jeewon R, Johnston PR, Kirk PM, Malosso E, May TW, Meyer W, Nilsson RH, Öpik M, Robert V, Stadler S, Thines M, Vu D, Yurkov AM, Zhang N, Schoch CL (2021) Fungal taxonomy and sequence-based nomenclature. Nature Microbiology 6(5): 540–548. https://doi.org/10.1038/s41564-021-00888-x

Marinov M, Fitzhugh K, Reis RE, Engel MS (2022) Putting the cart before the horse: SeqCode’s attempt to solve systematics issues with changes to nomenclature. Bionomina 31(1): 97–107. https://doi.org/10.11646/bionomina.31.1.6

May TW, Redhead SA, Lombard L, Rossman AY (2018) XI International Mycological Congress: report of Congress action on nomenclature proposals relating to fungi. IMA Fungus 9: xxii–xxvii. https://doi.org/10.1007/BF03449448

Murray RGE, Stackebrandt E (1995) Taxonomic note: implementation of the provisional status Candidatus for incompletely described prokaryotes. International Journal of Systematic and Evolutionary Bacteriology 45(1): 195–196. https://doi.org/10.1099/00207713-45-1-186

Nilsson RH, Ryberg M, Sjökvist E, Abarenkov K (2011) Rethinking taxon sampling in the light of environmental sequencing. Cladistics 27: 197–203. https://doi.org/10.1111/j.1096-0031.2010.00336.x

Nilsson RH, Wurzbacher C, Bahram M, Coimbra VRM, Larsson E, Tedersoo L, Eriksson J, Duarte C, Svantesson S, Sánchez-García M, Ryberg MK, Kristiansson E, Abarenkov K (2016) Top 50 most wanted fungi. MycoKeys 12: 29–40. https://doi.org/10.3897/mycokeys.12.7553

Nilsson RH, Larsson KH, Taylor AFS, Bengtsson-Palme J, Jeppesen TS, Schigel D, Kennedy P, Picard K, Glöckner FO, Tedersoo L, Saar I, Kõljalg U, Abarenkov K (2019) The UNITE database for molecular identification of fungi: handling dark taxa and parallel taxonomic classifications. Nucleic Acids Research 47(D1): D259–D264. https://doi.org/10.1093/nar/gky1022

R Core Team (2020). R: A language and environment for statistical computing. R Foundation for Statistical Computing, Vienna, Austria. URL https://www.R-project.org/

Retter A, Nilsson RH, Bourlat SJ (2019) Exploring the taxonomic composition of two fungal communities on the Swedish west coast through metabarcoding. Biodiversity Data Journal 7: e35332. https://doi.org/10.3897/BDJ.7.e35332

Ryberg M, Nilsson RH (2018) New light on names and naming of dark taxa. MycoKeys 30: 31–39. https://doi.org/10.3897%2Fmycokeys.30.24376

Tedersoo L, Bahram M, Puusepp R, Nilsson RH, James TY (2017) Novel soil-inhabiting clades fill gaps in the fungal tree of life. Microbiome 5(1): 1–10. https://doi.org/10.1186/s40168-017-0259-5

Tedersoo L, Anslan S, Bahram M, Kõljalg U, Abarenkov K (2020) Identifying the ‘unidentified’fungi: a global-scale long-read third-generation sequencing approach. Fungal Diversity 103: 273–293. https://doi.org/10.1007/s13225-020-00456-4

Tedersoo L, Mikryukov V, Zizka A, Bahram M, Hagh-Doust N, Anslan S, Prylutskyi O, Delgado-Baquerizo M, Maestre FT, Pärn J, Öpik M, Moora M, Zobel M, Espenberg M, Mander U, Khalid AN, Corrales A, Agan A, Vasco-Palacios A-V, Saitta A, Rinaldi AC, Verbeken A, Sulistyo BP, Tamgnoue B, Furneaux B, Ritter CD, Nyamukondiwa C, Sharp C, Marín C, Gohar D, Klavina D, Sharmah D, Dai DQ, Nouhra E, Biersma EM, Rähn E, Cameron EK, De Crop E, Otsing E, Davydov EA, Albornoz FE, Brearley FQ, Buegger F, Zahn G, Bonito G, Hiiesalu I, Barrio IC, Heilmann-Clausen J, Ankuda J, Kupagme JY, Maciá-Vicente JG, Fovo JD, Geml J, Alatalo JM, Alvarez-Manjarrez J, Põldmaa K, Runnel K, Adamson K, Bråthen KA, Pritsch K, Tchan KI, Armolaitis K, Hyde KD, Newsham KK, Panksep K, Lateef AA, Tiirmann L, Hansson L, Lamit LJ, Saba M, Tuomi M, Gryzenhout M, Bauters M, Piepenbring M, Wijayawardene N, Yorou NS, Kurina O, Mortimer PE, Meidl P, Kohout P, Nilsson RH, Puusepp R, Drenkhan R, Garibay-Orijel R, Godoy R, Alkahtani S, Rahimlou S, Dudov SV, Põlme S, Ghosh S, Mundra S, Ahmed T, Netherway T, Henkel TW, Roslin T, Nteziryayo V, Fedosov VE, Onipchenko VG, Yasanthika WAE, Lim YW, Soudzilovskaia NA, Antonelli A, Kõljalg U, Abarenkov K (2022) Global patterns in endemicity and vulnerability of soil fungi. Global Change Biology 28(22): 6696–6710. https://doi.org/10.1111/gcb.16398

Turland NJ, Wiersema JH, Barrie FR, Greuter W, Hawksworth DL, Herendeen PS, Knapp S, Kusber W-H, Li D-Z, Marhold K, May TW, McNeill J, Monro AM, Prado J, Price MJ, Smith GF (2018) International Code of Nomenclature for algae, fungi, and plants (Shenzhen Code) adopted by the Nineteenth International Botanical Congress Shenzhen, China, July 2017. Regnum Vegetabile 159. Glashütten: Koeltz Botanical Books. https://doi.org/10.12705/Code.2018

Zamora JC, Svensson M, Kirschner R, Olariaga I, Ryman S, Parra LA, Geml J, Rosling A, Adamčík S, Ahti T, Aime MC, Ainsworth AM, Albert L, Albertó E, Altés García A, Ageev D, Agerer R, Aguirre-Hudson B, Ammirati J, Andersson H, Angelini C, Antonín V, Aoki T, Aptroot A, Argaud D, Arguello Sosa BI, Aronsen A, Arup U, Asgari B, Assyov B, Atienza V, Bandini D, Baptista-Ferreira JL, Baral HO, Baroni T, Barreto RW, Beker H, Bell A, Bellanger JM, Bellù F, Bemmann M, Bendiksby M, Bendiksen E, Bendiksen K, Benedek L, Bérešová-Guttová A, Berger F, Berndt R, Bernicchia A, Biketova AY, Bizio E, Bjork C, Boekhout T, Boertmann D, Böhning T, Boittin F, Boluda CG, Boomsluiter MW, Borovička J, Brandrud TE, Braun U, Brodo I, Bulyonkova T, Burdsall Jr HH, Buyck B, Burgaz AR, Calatayud V, Callac P, Campo E, Candusso M, Capoen B, Carbó J, Carbone M, Castañeda-Ruiz RF, Castellano MA, Chen J, Clerc P, Consiglio G, Corriol G, Courtecuisse R, Crespo A, Cripps C, Crous PW, da Silva GA, da Silva M, Dam M, Dam N, Dämmrich F, Das K, Davies L, De Crop E, De Kesel A, De Lange R, De Madrignac Bonzi B, dela Cruz TEE, Delgat L, Demoulin V, Desjardin DE, Diederich P, Dima B, Dios MM, Divakar PK, Douanla-Meli C, Douglas B, Drechsler-Santos ER, Dyer PS, Eberhardt U, Ertz D, Esteve-Raventós F, Etayo Salazar JA, Evenson V, Eyssartier G, Farkas E, Favre A, Fedosova AG, Filippa M, Finy P, Flakus A, Fos S, Fournier J, Fraiture A, Franchi P, Franco Molano AE, Friebes G, Frisch A, Fryday A, Furci G, Galán Márquez R, Garbelotto M, García-Martín JM, García Otálora MA, García Sánchez D, Gardiennet A, Garnica S, Garrido Benavent I, Gates G, Gerlach ACL, Ghobad-Nejhad M, Gibertoni TB, Grebenc T, Greilhuber I, Grishkan B, Groenewald JZ, Grube M, Gruhn G, Gueidan C, Gulden G, Gusmão LFP, Hafellner J, Hairaud M, Halama M, Hallenberg N, Halling RE, Hansen K, Harder CB, Heilmann-Clausen J, Helleman S, Henriot A, Hernandez-Restrepo M, Herve R, Hobart C, Hoffmeister M, Høiland K, Holec J, Holien H, Hughes K, Hubka V, Huhtinen S, Ivančevi B, Jagers M, Jaklitsch W, Jansen AE, Jayawardena RS, Jeppesen TS, Jeppson M, Johnston P, Jørgensen PM, Kärnefelt I, Kalinina LB, Kantvilas G, Karadelev M, Kasuya T, Kautmanová I, Kerrigan RW, Kirchmair M, Kiyashko A, Knapp DG, Knudsen H, Knudsen K, Knutsson T, Kolařík M, Kõljalg U, Košuthová A, Koszka A, Kotiranta H, Kotkova V, Koukol O, Kout J, Kovács GM, Kříž M, Kruys Å, Kučera V, Kudzma L, Kuhar F, Kukwa M, Kumar TKA, Kunca V, Kušan I, Kuyper TW, Lado C, Læssøe T, Lainé P, Langer E, Larsson E, Larsson KH, Laursen G, Lechat C, Lee S, Lendemer JC, Levin L, Lindemann U, Lindström H, Liu X, Llarena Hernandez RC, Llop E, Locsmándi C, Lodge DJ, Loizides M, Lőkös L, Luangsa-ard J, Lüderitz M, Lumbsch T, Lutz M, Mahoney D, Malysheva E, Malysheva V, Manimohan P, Marin-Felix Y, Marques G, Martínez-Gil R, Marson G, Mata G, Matheny PB, Mathiassen GH, Matočec N, Mayrhofer H, Mehrabi M, Melo I, Meši A, Methven AS, Miettinen O, Millanes Romero AM, Miller AN, Mitchell JK, Moberg R, Moreau PA, Moreno G, Morozova O, Morte A, Muggia L, Muñoz González G, Myllys L, Nagy I, Nagy LG, Neves MA, Niemelä T, Nimis PL, Niveiro N, Noordeloos ME, Nordin A, Noumeur SR, Novozhilov Y, Nuytinck J, Ohenoja E, Oliveira Fiuza P, Orange A, Ordynets A, Ortiz-Santana B, Pacheco L, Pál-Fám F, Palacio M, Palice Z, Papp V, Pärtel K, Pawlowska J, Paz A, Peintner U, Pennycook S, Pereira OL, Pérez Daniëls P, Pérez-De-Gregorio Capella MÀ, Pérez del Amo CM, Pérez Gorjón S, Pérez-Ortega S, Pérez-Vargas I, Perry BA, Petersen JH, Petersen RH, Pfister DH, Phukhamsakda C, Piątek M, Piepenbring M, Pino-Bodas R, Pinzón Esquivel JP, Pirot P, Popov ES, Popoff O, Prieto Álvaro M, Printzen C, Psurtseva N, Purahong W, Quijada L, Rambold G, Ramírez NA, Raja H, Raspé O, Raymundo T, Réblová M, Rebriev YA, Reyes García JD, Ribes Ripoll MA, Richard F, Richardson MJ, Rico VJ, Robledo GL, Rodrigues Barbosa F, Rodriguez-Caycedo C, Rodriguez-Flakus P, Ronikier A, Rubio Casas L, Rusevska K, Saar G, Saar I, Salcedo I, Salcedo Martínez SM, Salvador Montoya CA, Sánchez Ramírez S, Sandoval-Sierra JV, Santamaria S, Santana Monteiro J, Schroers HJ, Schulz B, Schmidt-Stohn G, Schumacher T, Senn-Irlet B, Ševčíková H, Shchepin O, Shirouzu T, Shiryaev A, Siepe K, Sir EB, Sohrabi M, Soop K, Spirin V, Spribille T, Stadler M, Stalpers J, Stenroos S, Suija A, Sunhede S, Svantesson S, Svensson S, Svetasheva TYu, Świerkosz K, Tamm H, Taskin H, Taudière A, Tedebrand J-O, Tena Lahoz R, Temina M, Thell A, Thines M, Thor G, Thüs H, Tibell L, Tibell S, Timdal E, Tkalčec Z, Tønsberg T, Trichies G, Triebel D, Tsurykau A, Tulloss RE, Tuovinen V, Ulloa Sosa M, Urcelay C, Valade F, Valenzuela Garza R, van den Boom P, Van Vooren N, Vasco-Palacios AM, Vauras J, Velasco Santos JM, Vellinga E, Verbeken A, Vetlesen P, Vizzini A, Voglmayr H, Volobuev S, von Brackel W, Voronina E, Walther G, Watling R, Weber E, Wedin M, Weholt Ø, Westberg M, Yurchenko E, Zehnálek P, Zhang H, Zhurbenko MP, Ekman S (2018) Considerations and consequences of allowing DNA sequence data as types of fungal taxa. IMA Fungus 9(1): 167–175. https://doi.org/10.5598%2Fimafungus.2018.09.01.10

